# Cholesterol activates the G-protein coupled receptor Smoothened to promote morphogenetic signaling

**DOI:** 10.1101/070623

**Authors:** Giovanni Luchetti, Ria Sircar, Jennifer H. Kong, Sigrid Nachtergaele, Andreas Sagner, Eamon F. X. Byrne, Douglas F. Covey, Christian Siebold, Rajat Rohatgi

**Affiliations:** Departments of Biochemistry and Medicine, Stanford University School of Medicine, Stanford, California, United States of America; Division of Structural Biology, Wellcome Trust Centre for Human Genetics, University of Oxford, Oxford, UK; Department of Developmental Biology, Washington University School of Medicine, St. Louis, Missouri, United States of America; The Francis Crick Institute, Mill Hill Laboratory, London, UK

## Abstract

Cholesterol is necessary for the function of many G-protein coupled receptors (GPCRs). We find that cholesterol is not just necessary but also sufficient to activate signaling by the Hedgehog (Hh) pathway, a prominent cell-cell communication system in development. Cholesterol influences Hh signaling by directly activating Smoothened (SMO), an orphan GPCR that transmits the Hh signal across the membrane in all animals. Unlike most GPCRs, which are regulated by cholesterol through their heptahelical transmembrane domains, SMO is activated by cholesterol through its extracellular cysteine-rich domain (CRD). Residues shown to mediate cholesterol binding to the CRD in a recent structural analysis also dictate SMO activation, both in response to cholesterol and to native Hh ligands. Our results show that cholesterol can initiate signaling from the cell surface by engaging the extracellular domain of a GPCR and suggest that SMO activity may be regulated by local changes in cholesterol abundance or accessibility.

## Introduction

Cholesterol, which makes up nearly half of the lipid molecules in the plasma membrane of animal cells, can influence many signal transduction events at the cell surface. It plays an important role in modulating the function of cell-surface receptors, including G-protein coupled receptors (GPCRs), the largest class of receptors that transduce signals across the plasma membrane, and antigen receptors on immune cells (Burger et al., 2000; Pucadyil and Chattopadhyay, 2006; Swamy et al., 2016). The structures of several GPCRs reveal cholesterol molecules tightly associated with the heptahelical transmembrane domain (7TMD) (Cherezov et al., 2007; Ruprecht et al., 2004; Wu et al., 2014). Cholesterol can influence GPCR stability, oligomerization and ligand affinity (Fahrenholz et al., 1995; Gimpl et al., 1997; Gimpl and Fahrenholz, 2002; Prasanna et al., 2014; Pucadyil and Chattopadhyay, 2004). Cholesterol also organizes membrane microdomains, or “rafts,” containing proteins and lipids that function as platforms for the detection and propagation of extracellular signals (Lingwood and Simons, 2010). In all of these cases cholesterol plays a permissive role; however, it is not sufficient to trigger signaling on its own. Could cholesterol play a more instructive role— is it sufficient, not just necessary, to initiate signaling from the plasma membrane?

We find that cholesterol can indeed play an instructive signaling role in the Hedgehog (Hh) pathway, an iconic signaling system that plays roles in development, regeneration, and cancer. Multiple seemingly unrelated links have been described between cholesterol and Hh signaling (summarized in (Eaton, 2008; Incardona and Eaton, 2000)). While the best-defined role for cholesterol is in the biogenesis of Hh ligands (Porter et al., 1996), it also plays an independent role in the reception of Hh signals. Pharmacological or genetic depletion of cholesterol reduces cellular responses to Hh ligands, which has led to the view that cholesterol is *permissive* for Hh signaling (Blassberg et al., 2016; Cooper et al., 1998; Cooper et al., 2003; Incardona et al., 1998; Incardona and Roelink, 2000). Distinct from these previous observations, we find that an acute increase in plasma membrane cholesterol is *sufficient* to activate Hh signaling. Thus, cholesterol can initiate signals from the cell surface by acting as an activating ligand for a GPCR family protein.

## Results

### Cholesterol is sufficient to activate the Hedgehog signaling pathway

While testing the effect of a panel of sterol lipids on Hh signaling in cultured fibroblasts, we made the serendipitous observation that cholesterol could induce the transcription of Hh target genes. Since cholesterol is very poorly soluble in aqueous media, we delivered it to cultured cells as an inclusion complex (hereafter called MβCD:cholesterol) with the cyclic oligosaccharide Methyl-β−cyclodextrin (MβCD) (Zidovetzki and Levitan, 2007). Throughout this paper, we state the concentration of MβCD in the MβCD:cholesterol complexes, since this concentration is known exactly; for saturated complexes, the molar concentration of cholesterol is predicted to be ∼8-10-fold lower than that of MβCD (Christian et al., 1997; Klein et al., 1995). MβCD:cholesterol complexes have been shown to be the most effective way to rapidly increase cholesterol in the plasma membrane, the subcellular location for most transmembrane signaling events (Christian et al., 1997).

MβCD:cholesterol activated Hh signaling in NIH/3T3 cells and Mouse Embryonic Fibroblasts (MEFs), cultured cell lines that have been extensively used for mechanistic studies of the Hh pathway (Figure 1). MβCD:cholesterol treatment activated the transcription of *Gli1* (Figures 1A, 1B), a direct Hh target gene used as a measure of signal strength, and also reduced protein levels of the repressor form of the transcription factor GLI3, a consequence of signaling known to be independent of transcription (Figure 1B). MβCD:cholesterol induced a concentration-dependent, bell-shaped Hh signaling response (Figure 1A). Low doses of MβCD:cholesterol, which have only a minor effect on signaling, also increased the potency of the native ligand SHH, as seen by a leftward shift in the SHH dose-response curve (Figure 1C).

**Figure 1.**
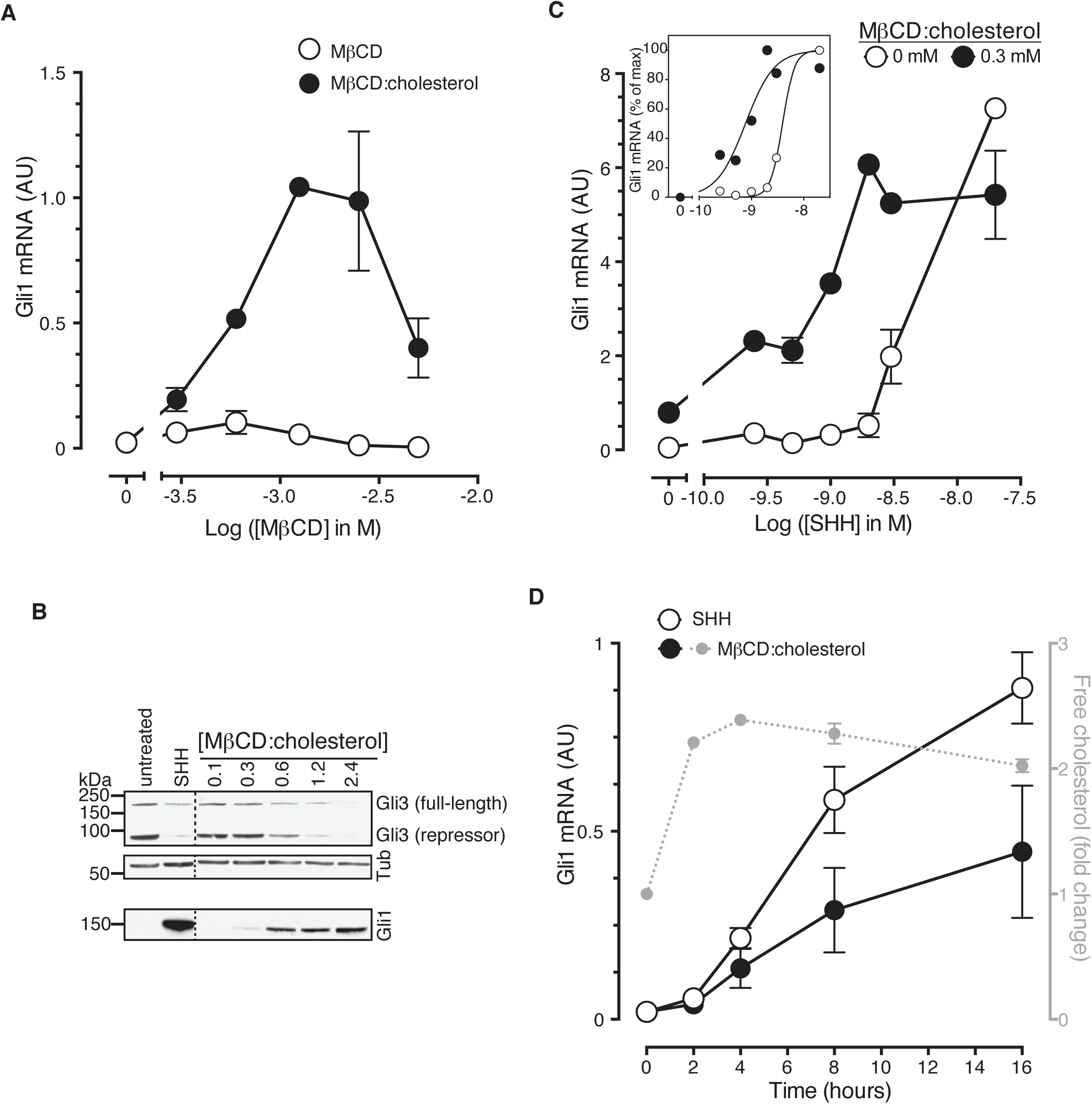
Cholesterol is sufficient to activate Hh target genes in NIH/3T3 cells. (**A**) *Gli1* mRNA, encoded by a direct Hh target gene, was measured by quantitative real-time reverse-transcription PCR (qRT-PCR) and normalized to mRNA levels of the housekeeping gene *GAPDH* after treatment (12 hours) with various doses of naked MβCD or a saturated MβCD:cholesterol (8.8:1 molar ratio) complex. In both cases, the concentration of MβCD is plotted on the abscissa. (**B**) Immunoblotting was used to measure protein levels of GLI1, full-length GLI3 and the repressor fragment of GLI3 after treatment (12 hours) with various concentrations (in mM) of MβCD:cholesterol. Dotted lines demarcate non-contiguous regions of the same immunoblot that were juxtaposed for clarity. (**C**) *Gli1* induction in response to various doses of SHH in the presence or absence of a low dose of MβCD:cholesterol. Inset shows non-linear curve fits to the data after a normalization in which the *Gli1* mRNA level in the absence of SHH was set to 0% and at the maximum dose of SHH was set to 100%. (**D**) Time course of *Gli1* induction (left y-axis) after treatment with SHH (265 nM) or the MβCD:cholesterol complex (2.5 mM). The gray circles (right y-axis) show the kinetics of increase in unesterified cholesterol (normalized to total protein) after the addition of MβCD:cholesterol. In all graphs, circles depict mean values from 3 replicates and error bars show the SD.

Cholesterol can influence multiple cellular processes at short and long timescales, so we compared the kinetics of MβCD:cholesterol-induced activation of *Gli1* to (1) the kinetics of MβCD:cholesterol-mediated delivery of cholesterol to cells and to (2) the kinetics of SHH-induced *Gli1* expression. Cholesterol loading of cells by MβCD:cholesterol was nearly complete by 2 hours, as determined by a standard enzymatic assay for free (unesterified) cholesterol (Figure 1D). The increase in cellular levels of free cholesterol was also confirmed by the transcriptional suppression of genes encoding enzymes in the pathway for cholesterol biosynthesis (Figure 1—Figure Supplement 1A). Importantly, there was a significant increase in the accessible or chemically active (Radhakrishnan and McConnell, 2000) pool of cholesterol in the plasma membrane, as shown by increased cell-surface labeling with a cholesterol-binding toxin (Perfringolysin O (PFO), Figure 1—Figure Supplement 1B)(Das et al., 2013). The initial activation of *Gli1* by MβCD:cholesterol coincided with the loading of cells with cholesterol, starting at 2 hours (Figure 1D). The kinetics of *Gli1* induction by MβCD:cholesterol paralleled those of *Gli1* induction by the native ligand SHH, despite the fact the absolute levels of signaling were higher in response to SHH. The rapid Hh signaling response to cholesterol, temporally correlated with the acute increase in cholesterol levels in the plasma membrane, is unlikely to be mediated by indirect or secondary transcriptional effects.

It was important to distinguish signaling effects caused by MβCD from those caused by cholesterol itself, especially because MβCD has been proposed to enhance Hh signaling by extracting an inhibitory sterol from cells (Sever et al., 2016). Following a previously-described protocol (Christian et al., 1997), we treated fibroblasts with a series of MβCD complexes in which the MβCD concentration was held constant at 1.25 mM while the cholesterol concentration was varied. Under these conditions, Hh signaling activity increased in proportion to the amount of cholesterol in the MβCD:cholesterol complexes (Figure 2A). Thus, cholesterol must be the active ingredient in these complexes that activates Hh signaling.

**Figure 2.**
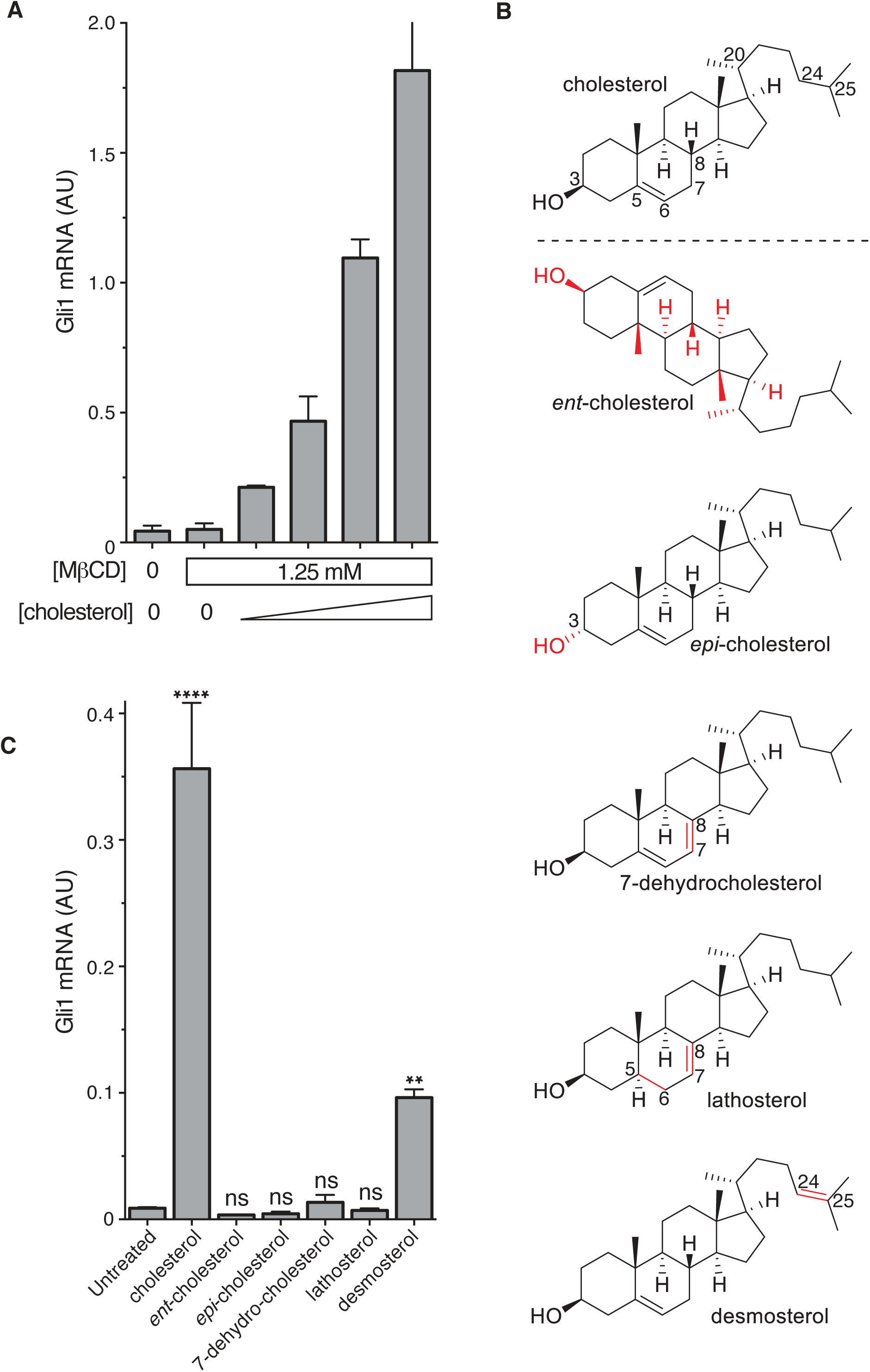
The cholesterol in MβCD:cholesterol complexes activates Hedgehog signaling. (**A**) Mean (+/−SD, n=3) *Gli1* mRNA levels after 12 hours of treatment of NIH/3T3 cells with a series of inclusion complexes in which the MβCD concentration was clamped at 1.25 mM while the cholesterol concentration was varied to yield MβCD:cholesterol molar ratios of 12:1, 9:1, 7:1 and 6:1. (**B**) Structures of cholesterol analogs tested for Hh signaling activity as inclusion complexes with MβCD. Structural differences from cholesterol are highlighted in red: *ent*-cholesterol is the mirror-image of cholesterol with inverted stereochemistry at all 8 stereocenters; *epi*-cholesterol is a diastereomer with inverted stereochemistry only at the 3 carbon postion; 7-dehydrocholesterol, lathosterol and desmosterol are naturally occurring cholesterol precursors. (**C**) Mean (+/−SD, n=4) *Gli1* mRNA levels after treatment (12 hours) with inclusion complexes of MβCD (1.25 mM) with the indicated sterols (see **B** for structures). Asterisks denote statistical significance for difference from the untreated sample using one-way ANOVA with a Holm-Sidak post-test.

To define the structural features of cholesterol required to activate Hh signaling, we used MβCD to deliver a panel of natural and synthetic analogs (Figure 2B). This experimental approach was inspired by previous studies of the cholesterol sensor SREBP cleavage-activating protein (SCAP)(Brown et al., 2002). The Hh signaling activity of cholesterol was exquisitely stereoselective— neither its enantiomer (*ent*-cholesterol) nor an epimer with an inverted configuration only at the 3-hydroxy position (*epi*-cholesterol) could activate Hh target genes (Figure 2C). Enantioselectivity is consistent with cholesterol acting through a chiral binding pocket on a protein target, rather than by altering membrane properties (Covey, 2009). Hh signaling activity was also lost when either the number or the position of double bonds in the tetracyclic sterol nucleus were altered in 7-dehydrocholesterol (7-DHC) and lathosterol, two endogenous biosynthetic precursors of cholesterol. Interestingly, desmosterol, another immediate biosynthetic precursor of cholesterol that contains an additional double-bond in the iso-octyl chain, retained signaling activity. This structure-activity relationship points to the tetracyclic ring, conserved between cholesterol and desmosterol, as the critical structural element required for activity. We cannot exclude the possibility that desmosterol activated signaling because it was rapidly converted to cholesterol in cells. These strict structural requirements suggest a specific, protein-mediated effect of cholesterol on the Hh signaling pathway and further exclude the possibility that signaling activity is due to extraction of an inhibitor from cells by MβCD (present at the same concentration in all the sterol complexes tested in Figure 2C).

MβCD:sterol inclusion complexes have been suggested to potentiate Hh signaling by depleting an inhibitory molecule through an exchange reaction (Sever et al., 2016). This model cannot explain our results because the concentration (Figure 2A) and structure (Figure 2C) of the sterol in the inclusion complex, despite an unchanging MβCD concentration, can modulate Hh signaling activity.

### Cholesterol functions at the level of Smoothened to activate Hedgehog signaling

A simplified schematic of the Hh signaling pathway is provided in Figure 3A (Briscoe and Therond, 2013). The receptor for Hh ligands, Patched 1 (PTCH1), inhibits signaling by suppressing the activity of SMO, a member of the GPCR superfamily. SHH binds and inhibits PTCH1, thereby allowing SMO to adopt an active conformation and transmit the Hh signal across the plasma membrane. Cytoplasmic signals from SMO overcome two negative regulators of the pathway, protein kinase A (PKA) and suppressor of fused (SUFU), ultimately leading to the activation and nuclear translocation of the GLI family of Hh transcription factors.

**Figure 3.**
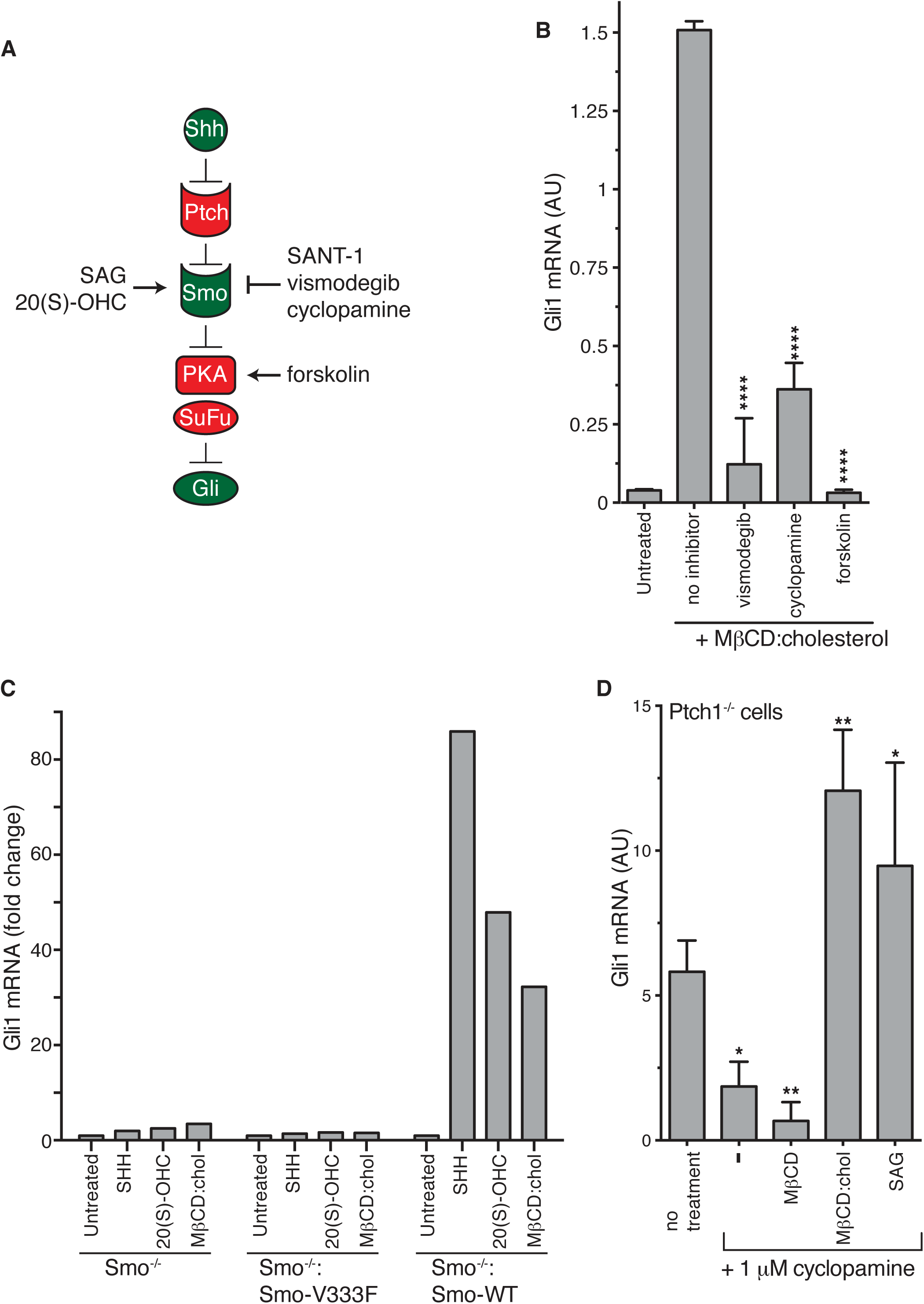
Smoothened activity is necessary for cholesterol to activate Hh signaling. (**A**) Schematic of the Hh signaling pathway showing the sequence in which core components function to transmit the signal from the cell surface to the nucleus. SAG and 20(S)-OHC are agonists and SANT-1, vismodegib, and cyclopamine are antagonists that bind and modulate the activity of SMO. Forskolin blocks signaling by elevating cAMP levels, which increases the activity of Protein Kinase A. (**B**) Mean (+/-SD, n=3) *Gli1* mRNA levels after treatment with MβCD:cholesterol (1.25 mM, 12 hours) in the presence of vismodegib (1 μM), cyclopamine (10 μM) or forskolin (10 μM). (**C**) Fold-change in *Gli1* mRNA levels after addition of agonists (12 hours) to Smo^−/−^ cells, in which both *Smo* alleles have been genetically inactivated, or Smo^−/−^ cells stably expressing a wild-type (WT) SMO protein or a variant SMO protein carrying an inactivating mutation (V333F) in its 7TMD (Byrne et al., 2016). SHH was used at 265nM, 20(S)-OHC at 5 μM, and MβCD:cholesterol at 1.25 mM. (**D**) Mean (+/− SD, n=4) *Gli1* mRNA levels in Ptch1^−/−^ cells after treatment with cyclopamine alone or cyclopamine in the presence of SAG (100 nM), MβCD (1.25 mM) or MβCD:cholesterol (1.25 mM). Asterisks denote statistical significance for difference from the “no inhibitor” sample in **B** and the “no treatment” sample in **D** using one-way ANOVA with a Holm-Sidak post-test.

To pinpoint the site of cholesterol action within this sequence of signaling events, we conducted a series of epistasis experiments (Figure 3). The addition of forskolin (Fsk), which leads to an increase in the activity of PKA, blocks Hh signaling at a step between SMO and the GLI proteins. Fsk inhibited MβCD:cholesterol-mediated signaling, placing the site of cholesterol action at the level of or upstream of PKA (Figure 3B). Two direct SMO antagonists, the steroidal natural product cyclopamine and the anti-cancer drug vismodegib, blocked *Gli1* activation by MβCD:cholesterol (Figure 3B)(Sharpe et al., 2015). This pharmacological profile established that MβCD:cholesterol requires SMO activity to promote signaling. Indeed, MEFs completely lacking SMO (*Smo*^−/−^ cells) failed to respond to MβCD:cholesterol, and the stable re-expression of wild-type (WT) SMO, but not a point mutant locked in an inactive conformation (Smo-V333F), rescued signaling (Figure 3C)(Varjosalo et al., 2006; Wang et al., 2014). Thus, cholesterol must activate the Hh pathway at the level of PTCH1, SMO or an intermediate step.

We evaluated the possibility that MβCD:cholesterol interferes with the function of PTCH1 by using *Ptch1*^−/−^ MEFs, which completely lack PTCH1 protein and have high levels of Hh target gene induction driven by constitutively activated SMO (Taipale et al., 2002). MβCD:cholesterol activated signaling in *Ptch1*^−/−^ cells treated with cyclopamine to partially suppress SMO activity, showing that cholesterol signaling activity did not depend on the presence of PTCH1 (Figure 3D). MβCD:cholesterol behaved much like the direct SMO agonist SAG, since both could overcome SMO inhibition by cyclopamine in the absence of PTCH1.

Our epistasis experiments pointed to SMO as the target of cholesterol. However, compared to treatment with the native ligand SHH, SMO did not accumulate to high levels in primary cilia in cells treated with MβCD:cholesterol (Figure 3—figure supplement 1A-1C), an observation that may explain the lower signaling efficacy of cholesterol compared to SHH.

### The cysteine-rich domain of Smoothened is required for the signaling activity of cholesterol

SMO contains two physically separable binding sites capable of interacting with steroidal ligands (Figure 4A) (Nachtergaele et al., 2012; Sharpe et al., 2015). Agonistic oxysterols, such as 20(S)-hydroxycholesterol (20(S)-OHC), engage a hydrophobic groove on the surface of the extracellular cysteine-rich domain (CRD) of SMO (Myers et al., 2013; Nachtergaele et al., 2013; Nedelcu et al., 2013). We recently reported that cholesterol could also occupy this CRD groove. A cholesterol molecule was resolved in this groove in a crystal structure of SMO. Furthermore, purified SMO bound to beads covalently coupled to cholesterol and this interaction could be blocked by free 20(S)-OHC, consistent with the view that both 20(S)-OHC and cholesterol occupy the same binding site (Byrne et al., 2016). In addition, the extracellular end of the SMO 7TMD binds to the steroidal alkaloid cyclopamine, as well as to several non-steroidal synthetic agonists and antagonists (Chen et al., 2002a; Chen et al., 2002b; Frank-Kamenetsky et al., 2002; Khaliullina et al., 2015).

**Figure 4.**
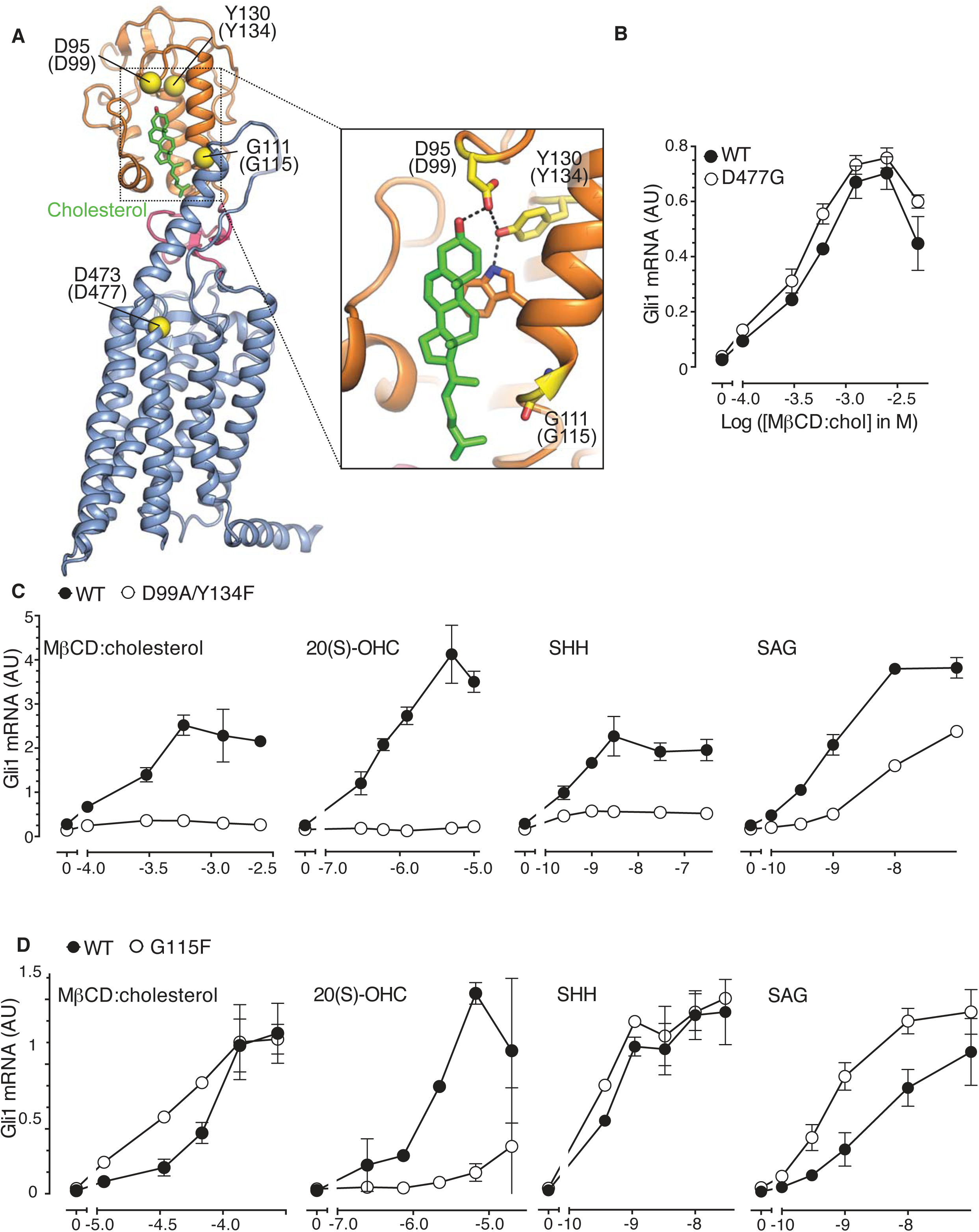
The SMO cysteine-rich domain is required for cholesterol-mediated activation of Hh signaling. (**A**) Structure of human SMO (PDB 5L7D), with the CRD in orange, the 7TMD in blue, the linker domain in pink, and the cholesterol ligand bound to the CRD in green. The Cα positions of the gatekeeper residues in the two ligand binding sites are highlighted as yellow spheres and numbered, with the mouse numbering shown in parenthesis. The inset shows a close-up of the cholesterol-binding site. D95 and Y130 form part of a hydrogen-bonding network (dotted lines) with the 3-hydroxyl of cholesterol, G111 abuts the iso-octyl chain of cholesterol, and D473 is a critical residue in the 7TMD binding-site. (**B, C** and **D**) Dose-response curves for the indicated agonists in Smo^−/−^ cells stably expressing WT SMO (always solid black circles) or the indicated SMO variants (open circles) carrying mutations in the 7TMD ligand-binding site (**B**) or at two opposite ends of the CRD binding groove (**C** and **D**). All agonists were applied to cells for 12 hours and mean (+/-SD) values for *Gli1* mRNA are plotted based on 3 replicates. In **C** and **D**, values on the abscissa represent Log([Agonist] in M) and the ordinate for all four graphs is only shown once at the left.

In order to distinguish if the activating effect of cholesterol is mediated by the cholesterol binding groove in the SMO CRD or the cyclopamine binding site in the 7TMD, we asked whether MβCD:cholesterol could activate signaling in *Smo*^−/−^ cells stably reconstituted with wild-type SMO (SMO-WT) or SMO variants carrying mutations in gatekeeper residues that have been shown to disrupt these two ligand-binding sites. The Asp477Gly mutation in the 7TM binding-site of SMO (Figure 4A), initially isolated from a patient whose tumor had become resistant to vismodegib, reduces binding and responsiveness to a subset of 7TM ligands, including SAG and vismodegib (Yauch et al., 2009). In the CRD, Asp99Ala/Tyr134Phe and Gly115Phe are mutations at opposite ends of the shallow sterol-binding groove that block the ability of 20(S)-OHC to both bind SMO and activate Hh signaling (Figure 4A)(Nachtergaele et al., 2013). The Asp99Ala and Tyr134Phe mutations disrupt a hydrogen-bonding network with the 3β-hydroxyl group of sterols (Figure 4A, inset)(Byrne et al., 2016).

The Asp477Gly mutation in the 7TMD domain had no effect on the ability of MβCD:cholesterol to activate Hh signaling (Figure 4B). SMO bearing a bulkier, charge-reversed mutation at this site (Asp477Arg) that increases constitutive signaling activity also remained responsive to MβCD:cholesterol (Figure 4—Figure Supplement 1A)(Dijkgraaf et al., 2011). In contrast, the Asp99Ala/Tyr134Phe mutation in the CRD reduced the ability of MβCD:cholesterol to activate Hh signaling (Figure 4C). The Asp99Ala/Tyr134Phe SMO mutant was also impaired in its responsiveness to SHH and to 20(S)-OHC, but remained responsive to the 7TMD ligand SAG (Figure 4C). A complete deletion of the CRD (SMO-ΔCRD), which increased basal SMO signaling activity like the Asp477Arg mutation, also abolished signaling responses to MβCD:cholesterol (Figure 4—Figure Supplement 1B)(Myers et al., 2013; Nedelcu et al., 2013). This mutational analysis supports the model that the CRD binding-site, rather than the 7TMD binding-site, mediates the effect of cholesterol on SMO activity and thus on Hh signaling.

Interestingly, a mutation in Gly115, which is located on the opposite end of the CRD ligand-binding groove (Figure 4A), did not alter the response to MβCD:cholesterol, even though it diminished the response to 20(S)-OHC as previously noted (Figure 4D)(Nachtergaele et al., 2013). The SMO-Gly115Phe mutant also responded normally to the native ligand SHH (Figure 4D). Gly115 is located near the iso-octyl chain of cholesterol in the SMO structure (Figure 4A). The introduction of a bulky, hydrophobic phenyl group at residue 115 may prevent the hydroxyl in the iso-octyl chain of 20(S)-OHC from being accommodated in the binding groove, but not disrupt binding of the purely hydrophobic iso-octyl chain of cholesterol. The ability of mutations to segregate 20(S)-OHC responses from cholesterol responses is consistent with solution-state small-angle X-Ray scattering data showing distinct conformations for SMO bound to these two steroidal ligands (Byrne et al., 2016).

The ability of the Gly115Phe mutation to distinguish between cholesterol and 20(S)-OHC responses allowed us to address an important outstanding question: could cholesterol activate SMO only after being oxidized to a side-chain oxysterol? In addition to 20(S)-OHC, oxysterols carrying hydroxyl groups on the 25 and 27 positions can bind and activate SMO (Corcoran and Scott, 2006; Dwyer et al., 2007; Myers et al., 2013; Nachtergaele et al., 2012). However, 20(S)-OHC, 25-OHC and 27-OHC, when delivered to cells as MβCD conjugates, were all significantly compromised in their ability to activate Hh signaling in cells expressing SMO-Gly115Phe (Figure 4—Figure Supplement 1C). In contrast, cholesterol-induced signaling was unaffected (Figure 4—Figure Supplement 1D); therefore, cholesterol must not be activating signaling by being metabolized to one of these side-chain oxysterols. Instead, our data suggests that cholesterol can directly activate Hh signaling through the CRD of SMO.

### Cholesterol can drive the differentiation of spinal cord progenitors

Our mechanistic experiments in cultured fibroblasts led us to ask whether cholesterol could also promote Hh-dependent cell differentiation decisions. In the developing vertebrate spinal cord, the Hh ligand Sonic Hedgehog (SHH) acts as a morphogen to specify the dorsal-ventral pattern of progenitor subtypes (Figure 5A)(Jessell, 2000). This spatial patterning process can be recapitulated *in vitro*. Mouse neural progenitors exposed to increasing concentrations of SHH will express transcription factors that mark differentiation towards progressively more ventral neural subtypes: low, medium and high Hh signaling will generate progenitor subtypes positive for Nkx6.1, Olig2, and Nkx2.2, respectively (Dessaud et al., 2008; Gouti et al., 2014; Kutejova et al., 2016).

**Figure 5.**
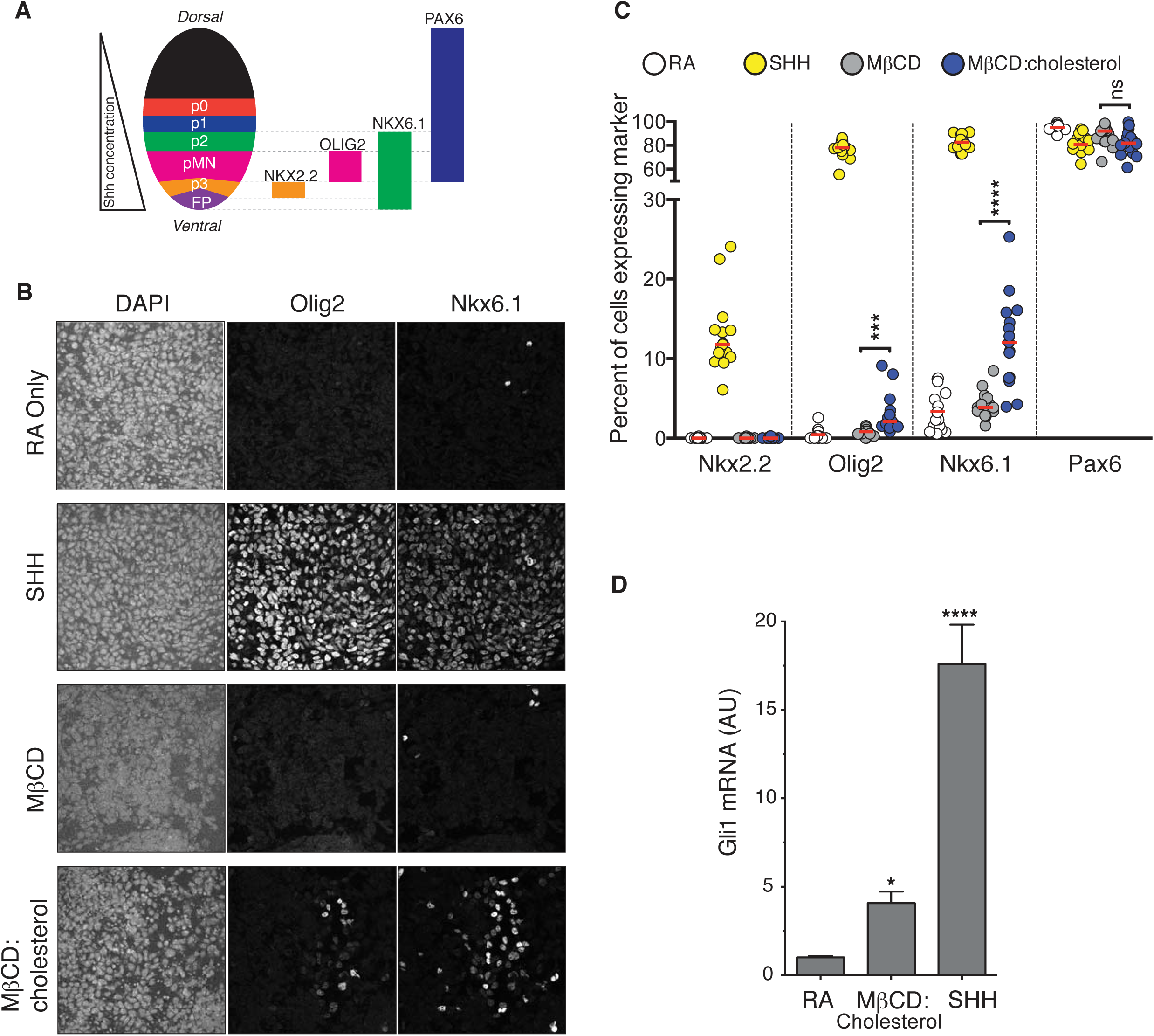
Cholesterol induces the differentiation of neural progenitors. (**A**) A schematic illustrating the relationship between marker proteins used to assess differentiation and progenitor cell populations in the embryonic neural tube (taken from (Niewiadomski et al., 2014)). pFP – floor plate progenitors, pMN – motor neuron progenitors, p0, p1, p2, p3 – ventral interneuron progenitors. **(B)** Differentiation of neural progenitors was assessed by immunostaining for Nkx6.1+ and Olig2+ expression (see **A**) after treatment (48 hours) with Retinoic Acid (RA, 100 nM) alone or RA plus SHH (25 nM), MβCD (2 mM) or the saturated MβCD:cholesterol inclusion complex (2 mM). The percentage of nuclei (stained with DAPI) positive for four differentiation markers (see **A**) in 15 different images is plotted in (**C**), with each point representing one image of the type shown in (**B**) and the red line drawn at the median value. Asterisks denote statistical significance (unpaired *t*-test, Holm-Sidak correction, n=15) for the comparison between cells treated with RA+MβCD and RA+MβCD:cholesterol. (**D**) *Gli1* mRNA (mean +/− SD, n=3) after 48 hours of the indicated treatments. Asterisks denote statistical significance for difference from the RA-treated sample using one-way ANOVA with a Holm-Sidak post-test.

MβCD:cholesterol induced the formation of both Nkx6.1^+^ and Olig2^+^ progenitor subtypes at a low frequency in cultures of mouse spinal cord progenitors (Figures 5B, 5C) and also activated the transcription of *Gli1* (Figure 5D). The efficacy of both *Gli1* induction and ventral neural specification induced by MβCD:cholesterol were significantly less than those produced by a saturating concentration of SHH. However, we note that MβCD:cholesterol inclusion complexes could not be delivered at higher concentrations due to deleterious effects on the adhesion and viability of neural progenitors. Taken together, these observations suggest that MβCD:cholesterol is sufficient to activate low-level Hh signals in neural progenitors and consequently to direct differentiation towards neural cell types that depend on such signals.

## Discussion

To establish a causal or regulatory role for a component in a biological pathway, experiments should demonstrate that the component is both *necessary* and *sufficient* for activity. Cholesterol has been shown to be necessary for SMO activation, based on experiments using inhibitors of cholesterol biosynthesis and high concentrations (∼10 mM) of naked MβCD to strip the plasma membrane of cholesterol (Cooper et al., 2003). Impaired SMO activation caused by cholesterol deficiency has also been noted in Smith-Lemli-Opitz syndrome (SLOS), a congenital malformation syndrome caused by defects in the enzyme that converts 7-dehydrocholesterol to cholesterol (Blassberg et al., 2016; Cooper et al., 2003). In contrast to our results, the SMO CRD is dispensable for this permissive role of cholesterol. The depletion of cholesterol reduces signaling by SMO mutants lacking the entire CRD (Myers et al., 2013) or carrying mutations in the CRD binding-groove (Blassberg et al., 2016). By analogy with other GPCRs, these permissive effects are likely to be mediated by the SMO 7TMD.

We now find that cholesterol is also sufficient to activate Hh signalling in a dose-dependent manner. This instructive effect is mediated by the Class F GPCR SMO and maps to its extracellular CRD. Cholesterol engages a hydrophobic groove on the surface of the CRD, a groove that was previously shown to mediate the activating influence of oxysterols (Myers et al., 2013; Nachtergaele et al., 2013; Nedelcu et al., 2013) and represents an evolutionarily conserved mechanism for detecting hydrophobic small-molecule ligands (Bazan and de Sauvage, 2009). An analogous mechanism is present in the Frizzled family of Wnt receptors, where the Frizzled CRD binds to the palmitoleyl group of Wnt ligands, an interaction that is required for Wnt signaling (Janda et al., 2012). Thus, the instructive effects of cholesterol revealed in our present study and the permissive effects of cholesterol reported previously map to distinct, separable SMO domains.

There are many reasons why this activating effect of cholesterol on Hh signalling may not have been appreciated previously despite the fact that the activating effects of side-chain oxysterols have been known for a decade (Corcoran and Scott, 2006; Dwyer et al., 2007). First, the method of delivery, as an inclusion complex with MβCD, is critical to presenting cholesterol, a profoundly hydrophobic and insoluble lipid, in a bioavailable form capable of activating Smo. Even clear solutions of cholesterol in the absence of carriers like MβCD contain microcrystalline deposits or stable micelles that sequester cholesterol (Haberland and Reynolds, 1973). In contrast, side-chain oxysterols, which harbor an additional hydroxyl group, are significantly more hydrophilic and soluble in aqueous solutions, shown by their ∼50-fold faster transfer rates between membranes (Theunissen et al., 1986). Second, cholesterol levels in the cell are difficult to manipulate because they are tightly controlled by elaborate homeostatic signalling mechanisms (Brown and Goldstein, 2009). MβCD:cholesterol inclusion complexes have been shown to be unique in their ability to increase the cholesterol content of the plasma membrane rapidly at timescales (∼1-4 hours) at which cytoplasmic signaling pathways operate (Christian et al., 1997; Yancey et al., 1996). Other methods of delivery using low density lipoprotein particles and lipid dispersions, or mutations in genes regulating cholesterol homeostasis, function on a much slower time scale and are thus more likely to be confounded by indirect effects given the myriad cellular processes affected by cholesterol (Christian et al., 1997). Finally, the bell-shaped Hh signal-response curve (Figure 1A) implies that MβCD:cholesterol must be delivered in a relatively narrow, intermediate concentration range (1-2 mM) to observe optimal activity, with higher (>5 mM) concentrations commonly used to load cells with cholesterol producing markedly lower levels of signaling activity.

Our results are particularly informative in light of the recently solved crystal structure of SMO, unexpectedly found to contain a cholesterol ligand in its CRD groove (Figure 4A) (Byrne et al., 2016). Molecular dynamics simulations showed that cholesterol can stabilize the extracellular domains of SMO (Byrne et al., 2016), but the function of this bound cholesterol, whether it is an agonist, antagonist or co-factor, remains an important unresolved question in SMO regulation. Structure-guided point mutations in CRD residues that form hydrogen-bonding interactions with the 3β-hydroxyl of cholesterol, reduced signaling by cholesterol (Figure 4C) making it likely that cholesterol activates SMO by binding to the CRD in the pose revealed in the structure (Figure 4A). Thus, the cholesterol-bound SMO structure may very well represent an active-state conformation of the CRD.

A surprising feature of the structure is that CRD-bound cholesterol is located at a considerable distance (∼12 Å) away from the membrane, which would require a cholesterol molecule to desolvate from the membrane and become exposed to water in order to access its CRD binding pocket (Byrne et al., 2016) (Figure 6). The kinetic barrier, or the activation energy (Δ*G*^‡^), for this transfer reaction is predicted to be high (∼20 kcal/mole), based on the Δ*G*^‡^ for cholesterol transfer between two acceptors through an aqueous environment (Yancey et al., 1996). The unique ability of MβCD to shield cholesterol from water while allowing its rapid transfer to acceptors would allow it to bypass this kinetically unfavorable step by delivering it to the CRD binding site (Figure 6). These considerations present a regulatory puzzle for future research: how does cholesterol gain access to the CRD-binding pocket without MβCD and is this process regulated by native Hh ligands? Indeed, the kinetic barrier for cholesterol transfer to the CRD pocket makes it an ideal candidate for a rate-limiting, regulated step controlling SMO activity in cells.

**Figure 6.**
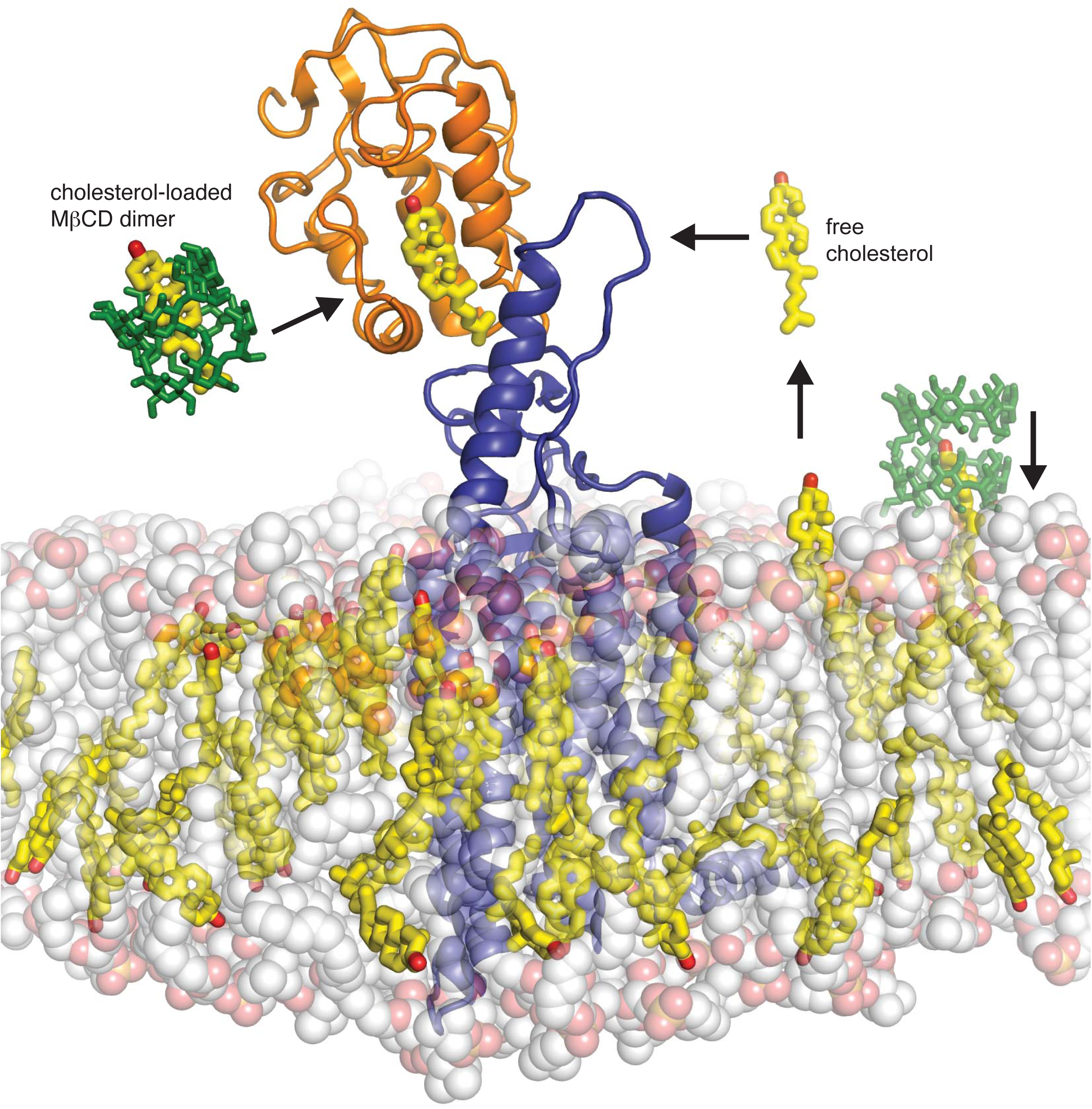
Models for how cholesterol may gain access to its binding-site in the SMO cysteine-rich domain. The structure of SMO bound to cholesterol (PDB 5L7D) is shown embedded in a lipid bilayer composed of 1-palmitoyl-2-oleoyl-sn-glycero-3-phosphocholine (POPC) and cholesterol in a ratio of 3:1 (Byrne et al., 2016). The SMO CRD is colored orange and the 7TMD is colored blue. Two molecules of MβCD (PDB QKH, shown as green sticks) form an inclusion complex with each molecule of cholesterol (PDB CLR, colored yellow in stick representation with the 3-hydroxyl shown red). MβCD could deliver cholesterol directly to the CRD binding pocket (left) or to the outer leaflet of the plasma membrane (right), which would subsequently require a second transfer step from the membrane to the CRD. The activation energy for the direct delivery mechanism on the left (<10 kcal/mole) is much lower than for the mechanism on the right (∼20 kcal/mole), where cholesterol has to desolvate from the membrane without a carrier to access the CRD site (Lopez et al., 2011; Yancey et al., 1996).

MβCD:cholesterol was consistently less active than the native ligand SHH in our assays (Figures 1D, 5C and 5D). Comparing the doses of MβCD:cholesterol to the doses of SHH delivered to cells is difficult. SHH was used at saturating concentrations; however, we could not assess the effects of MβCD:cholesterol at saturating doses, because the downward phase of the bell-shaped dose-response curve (in cultured fibroblasts, Figure 1A) and cell toxicity (in neural progenitors) proved to be dose-limiting. Aside from these technical considerations related to delivery, other possibilities for lower activity include the observation that MβCD:cholesterol did not induce the high-level accumulation of SMO in primary cilia (Figure 3—figure supplement 1) and the possibility that a different ligand regulates high-level signaling by SMO. Mutations in the 7TMD binding-site do not alter the constitutive or SHH-induced signaling activity of SMO, which has led to view that this site does not regulate physiological signaling (Myers et al., 2013; Yauch et al., 2009). In contrast, mutations in the cholesterol-binding site impaired responses to SHH (Byrne et al., 2016). Hence, a putative alternate ligand would have to engage a third, undefined site. Lastly, the presence of active PTCH1 is a major difference between SHH- and MβCD:cholesterol-induced signaling. The biochemical activity of PTCH1 (which is inactivated by SHH) may oppose the effects of MβCD:cholesterol, limiting signaling responses. Interestingly, MβCD:cholesterol was able to restore maximal Hh responses in the absence of PTCH1 (Figure 3D).

Our results may have implications for understanding how PTCH1 inhibits SMO, a longstanding mystery in Hh signaling. The necessity and sufficiency of cholesterol for SMO activation, mediated through two different regions of the molecule, means that SMO activity is likely to be highly sensitive to both the abundance and the accessibility of cholesterol in its membrane environment. Furthermore, PTCH1 has homology to a lysosomal cholesterol transporter, the Niemann-Pick C1 (NPC1) protein (Carstea et al., 1997), and PTCH1 has been purported to have cholesterol binding and transport activity (Bidet et al., 2011). Thus, our work supports a model where PTCH1 may inhibit SMO by reducing cholesterol content or cholesterol accessibility (or chemical activity) in a localized membrane compartment (such as the base of primary cilia) that contains SMO, leading to alterations in SMO conformation or trafficking (Bidet et al., 2011; Incardona et al., 2002; Khaliullina et al., 2009). Further tests of this hypothesis will require analysis of the biochemical activities of purified SMO and PTCH1 reconstituted into cholesterol-containing membranes. While cholesterol is an abundant lipid, clearly critical for maintaining membrane biophysical properties and for stabilizing membrane proteins, our work suggests that it may be also used as a second messenger to instruct signaling events at the cell surface through GPCRs and perhaps other cell-surface receptors.

## Acknowledgements

We thank Andres Lebensohn, Ganesh Pusapati, James Briscoe, Robert Blassberg and George Hedger for discussions and comments on the manuscript. This work was supported by the US National Institutes of Health (GM106078 and HL067773), Cancer Research UK (C20724/A14414), and the Taylor Family Institute for Psychiatric Research. G.L. was supported by the Ford Foundation, S.N. by the National Science Foundation and E.F.X.B. by NDM Oxford. A.S. has received funding from an EMBO LTF (1438-2013), HFSP LTF (LT000401/2014-L) and the People Programme (Marie Curie Actions) of the European Union's Seventh Framework Programme FP7-2013 under REA grant agreement n° 624973.

## Materials and Methods

### Cells and Reagents

### Reagents and Cell Lines

NIH/3T3 and 293 T cells were obtained from ATCC, *Smo*^−/−^ fibroblasts have been described previously (Varjosalo et al., 2006) and were originally obtained from Drs. James Chen and Philip Beachy. Suppliers for chemicals included Enzo Life Sciences (SAG), Toronto Research Chemicals (Cyclopamine), from, EMD Millipore (SANT-1), Tocris (20(S)-OHC), LC Labs (Vismodegib), Steraloids (25-OHC, 26-OHC, *epi*-cholesterol), Sigma (cholesterol, desmosterol, lathosterol, 7-dehydrocholesterol, Methyl-β-cyclodextrin), and Thermo Fisher (Alexa Fluor 647 NHS ester). *Ent*-cholesterol was synthesized as described previously (Jiang and Covey, 2002). Antibodies against GLI3 and GLI1 were from R&D Systems (AF3690) and Cell Signaling Technologies (Cat#L42B10) respectively. Human SHH carrying two isoleucine residues at the N-terminus and a hexahistidine tag at the C-terminus was expressed in *Escherichia coli* Rosetta(DE3)pLysS cells and purified by immobilized metal-affinity chromatography followed by gel-filtration chromatography as described previously (Bishop et al., 2009). Perfringolysin O (PFO) was purified as previously described (Das et al., 2013; Li et al., 2015) and covalently labeled with Alexa Fluor 647 dye following the manufacturer's instructions (Thermo Fisher Scientific).

### Methyl-β-Cyclodextrin Sterol Complexes

Sterols were dissolved in a mixture of chloroform-methanol (2:1 vol/vol) to generate a 10 mg / mL stock solution. To a glass vial, 8.7 μmole of sterol was delivered from the organic stock solution. Nitrogen gas was streamed over the sterol solution until the organic solvent was evaporated completely, generating a thin film in the vial. MβCD was dissolved in Opti-MEM at a final concentration of 50 mg / mL (38 mM), and 2 mL of this solution was added to the dried sterol film in the glass vial. A micro-tip sonicator was used to dissolve the mixture until it became clear. Solutions were filtered through a 0.1 μm filter and stored in glass vials at 4°C. Unless otherwise stated, the MβCD:cholesterol ratio was 8.8:1 in inclusion complexes. Preparation of the different ratios of cholesterol to MβCD (Figure 2) was achieved following the aforementioned protocol changing only the initial molar amount of cholesterol keeping the molar concentration of MβCD constant.

### Constructs

Constructs encoding mutant mouse SMO (D99A/Y134F, G115F, V333F, D477G, D477R, D477R/Y134F) were generated using the QuikChange method in the pCS2+:mSmo vector (Byrne et al., 2016)and then transferred by Gibson cloning to a retroviral vector (pMSCVpuro) for stable cell line construction.

### Stable cell lines

Stable cell lines were prepared as described previously by infecting Smo^−/−^ mouse embryonic fibroblasts with a retrovirus carrying untagged Smo variants cloned into pMSCVpuro (Byrne et al., 2016; Rohatgi et al., 2009). Retroviral supernatants were produced after transient transfection of Bosc23 helper cells with the pMSCV constructs (Pear et al., 1993). Virus-containing media from these transfections was directly used to infect Smo^−/−^ fibroblasts, and stable integrants were selected with puromycin (2 µg/mL). Cell lines stably expressing SMO-D99A/Y134F, SMO-V333F, SMO-D477R, and SMO-ΔCRD have been described and characterized previously, including measurement of SMO protein levels by immunoblotting (Byrne et al., 2016).

### Hedgehog signaling assays using quantitative RT-PCR

Stable cell lines expressing SMO variants or NIH/3T3 cells were grown to confluency in Dulbecco's Modified Eagle's Medium (DMEM) containing 10% Fetal Bovine Serum (FBS, Optima Grade, Atlanta Biologicals). Confluent cells were exchanged into 0.5% FBS DMEM for 24 hours to allow ciliogenesis prior to treatment with drugs and/or ligands in DMEM containing 0.5% FBS for various times, as indicated in the figure legends. The mRNA levels of *Gli1*, a direct Hh target gene commonly used as a metric for signalling strength, were measured using the *Power SYBR Green Cells-To-CT* kit (Thermo Fisher Scientific). The primers used are *Gli1* (forward primer: 5’-ccaagccaactttatgtcaggg-3’ and reverse primer: 5’-agcccgcttctttgttaatttga-3’), *Gapdh* (forward primer: 5’-agtggcaaagtggagatt-3’ and reverse primer: 5’-gtggagtcatactggaaca-3’), *Hmgcr* (forward primer: 5’-tgtggtttgtgaagccgtcat-3’ and reverse primer: 5’-tcaaccatagcttccgtagttgtc-3’), and *Hmgcs1* (forward primer: 5’-gggccaaacgctcctctaat-3’ and reverse primer: 5’-agtcataggcatgctgcatgtg-3’). Transcript levels relative to Gapdh were calculated using the ΔCt method. Each qRT-PCR experiment, which was repeated 3-4 times, included two biological replicates, each with two technical replicates.

### Data Analysis

Each experiment shown in the paper was repeated at least three independent times with similar results. All data was analyzed using GraphPad Prism. All points reflect mean values calculated from at least 3 replicates and error bars denote standard deviation (SD). The statistical tests used to evaluate significance are noted in the figure legends. Statistical significance in the figures is denoted as follows: ns: p>0.05, *: p≤0.05, **p≤0.01, ***p≤0.001, ****p≤0.0001.

### Mouse Embryonic Stem Cell Culture and Cell Differentiation

For maintenance, MM13 mouse embryonic stem cells (mESCs)(Wichterle et al., 2002) were plated on irradiated primary mouse embryonic fibroblasts (pMEFs) and cultured in mESC media (Dulbecco's Modified Eagle's Medium high glucose (Hyclone) and 15% Optima FBS (Atlanta Biologicals) supplemented with 1% MEM non-essential amino acids (Gibco), 1% penicillin/streptomycin (Gemini Bio-Products), 2 mM L-glutamine (Gemini Bio-Products), 1% EmbryoMax nucleosides (Millipore), 55 µM 2-mercaptoethanol (Gibco), and 1000 U/ml ESGRO LIF (Millipore). The mESCs were differentiated as previously described with minor modifications (Gouti et al., 2014; Ying et al., 2003). Briefly, the pMEFs were removed from the mESCs by dissociating the cells with 0.25% Trypsin/EDTA and then incubating the cells on tissue culture plates for two short successive periods (20 min each). To induce differentiation, the cells were plated on Matrigel (BD Biosciences) coated glass coverslips (12 mm diameter, placed in a 24-well plate) at a density of 2.4×10^4^ cells per coverslip in N2B27 media (Dulbecco's Modified Eagle's Medium F12 (Gibco) and Neurobasal Medium (Gibco) (1:1 ratio) supplemented with N-2 Supplement (Gibco), B-27 Supplement (Gibco), 1% penicillin/streptomycin (Gemini Bio-Products), 2 mM L-glutamine (Gemini Bio-Products), 40 µg/ml Bovine Serum Albumin (Sigma), and 55 µM 2-mercaptoethanol (Gibco)). On Day 0 and Day 1, cells were cultured in N2B27 with 10 ng/ml bFGF (R&D). On Day 2, the media was changed and the cells were cultured in N2B27 with 10 ng/ml bFGF (R&D) and 5 µM CHIR99021 (Axon). On Day 3, the media was changed and the cells were cultured in 1 ml of N2B27 supplemented with 100 nM Retinoic Acid (RA), 100 nM RA and 25 nM SHH, 100 nM RA and 2 mM MeβCD, or 100 nM RA and 2 mM MβCD+0.23 mM cholesterol. On Day 4, 1 ml N2B27 with 100 nM RA was added to each well, thus diluting each treatment condition by half. On Day 5 the cells were rinsed and fixed for further analysis.

### Immunofluorescence

NIH/3T3 cells were cultured in Dulbecco's modified Eagle's Medium (DMEM) containing 10% Fetal Bovine Serum (FBS, Optima Grade, Atlanta Biologicals) in 24-well plates at an initial density of 7.5×10^4^ on acid-washed glass cover-slips that were pre-coated with poly-L-lysine. Confluent cells were exchanged into 0.5% FBS DMEM to induce ciliogenesis for 24 hours. Ciliated cells were treated with the indicated drugs each dissolved in 0.5% FBS DMEM. Samples were fixed using 4% paraformaldehyde in phosphate buffered saline (PBS) for 10 minutes and washed three times with PBS. For SMO localization studies, cells were blocked and permeabilized in 1% donkey serum, 10 mg / mL bovine serum albumin (BSA), 0.1% triton X-100, and PBS. Primary antibodies were administered in block buffer for 2 hours at room temperature. Cover-slips were washed three times with wash buffer containing PBS and 0.1% triton X-100. Secondary antibodies were administered in block buffer for 1 hour. Cover-slips were washed three more times in wash buffer and mounted on glass slides using Pro-Long Diamond Antifade Mountant with DAPI (Life Technologies). For PFO staining, cells were fixed in 4% PFA, washed three times with PBS and stained with PFO in PBS without detergent. Cover-slips were washed three times with PBS and mounted on glass slides using Pro-Long Diamond Antifade Mountant with DAPI (Life Technologies). Images were acquired on an inverted Leica SP8 laser scanning confocal microscope with a 63X oil immersion objective (NA 1.4) using a HyD hybrid detector. Z-stacks were acquired with identical acquisition settings (gain, offset, laser power, frame format) within a given experiment. The following primary antibodies were used: rabbit anti-Smo (1:500)(Rohatgi et al., 2007), guinea pig anti-Arl13b(Pusapati et al., 2014), goat anti-GFP (1:2000) (Rockland), mouse anti-Nkx2.2 (1:100) (74.5A5, Developmental Studies Hybridoma Bank), mouse anti-Nkx6.1 (1:100) (F55A10, Developmental Studies Hybridoma Bank), guinea pig anti-Olig2 (1:20,000) (Novitch et al., 2001), rabbit anti-Pax6 (1:1000) (AB2237, Millipore). The following secondary antibodies were used: Alexa Fluor 488, Alexa Fluor 594, and Alexa Fluor 647 (Thermo Fisher Scientific).

### Image Analysis

Image processing for ciliary SMO levels was carried out using maximum projection images of the acquired Z-stacks using ImageJ. For quantification of ciliary Smo, first a mask was constructed using the Arl13b image (primary cilia marker), and then the mask was applied to the corresponding Smo image where the integrated fluorescence was measured. An identical region outside the cilia was measured to determine background fluorescence. Background correction was applied on a per cilia basis by subtracting the background fluorescence from the cilia fluorescence.

For neural differentiation experiments, fluorescent images were collected on a Leica TCS SP8 confocal imaging system equipped with a 40x oil immersion objective using the Leica Application Suite X (LASX) software. For each experiment, coverslips from each condition were grown, collected, and processed together to ensure that the cells were fixed and stained for the same duration of time. To ensure uniformity in imaging, the gain, offset, and laser power settings on the microscope were held constant for each antibody. At least 15 images were taken per condition. To ensure all cells were represented, z-stacks were acquired and counts were performed on the compressed images. Cell counts were conducted using the NIH ImageJ software suite with cell counter plugin. In total, 5000-6000 cells were analyzed per condition. The experiment was conducted independently a total of three times. Representative images shown in Figure 5 were processed equally using Adobe Photoshop, Adobe illustrator, and CorelDraw software.

### Cholesterol Quantification

Cells were cultured in Dulbecco's modified Eagle's Medium (DMEM) containing 10% Fetal Bovine Serum (FBS, Optima Grade, Atlanta Biologicals) in 6-well plates at an initial density of 3×10^5^ cells / well. Confluent cells were switched into 0.5% FBS DMEM to induce ciliogenesis for 24 hr. Cells were treated with indicated drugs dissolved in 0.5% FBS DMEM in duplicate. One sample was used to measure total protein by bicinchoninic acid assay (BCA), and the second for total lipid extraction and subsequent cholesterol quantification. Cells were washed once with Phosphate Buffered Saline (PBS), and harvested using a Corning cell lifter in PBS. The cell suspension was transferred to a 1.5 mL ependorf tube, centrifuged at 1000x g and the PBS aspirated. Total lipids were extracted from the cell pellet by the addition of 200 μL of chloroform-methanol (2:1 vol/vol). To induce phase separation, 100 μL of PBS was added to the lipid extract and the sample was centrifuged at 5,000x g for 5 minutes. The organic layer was transferred to a fresh 1.5 mL ependorf tube and the solvent removed under reduced pressure. Relative total free cholesterol was measured using the *Amplex Red Cholesterol Assay Kit* (Thermo Fisher Scientific) following the manufacturer's instructions. Lysis buffer containing 50 mM Tris pH 7.4, 150 mM NaCl, 2% Nonidet P-40, 0.5% sodium deoxycholate, 0.1% sodium dodecyl sulfate, 1 mM dithithreitol, and Sigma Fast protease inhibitor cocktail (Sigma-Aldrich) was used to disrupt the cell pellet. A ratio of total free cholesterol to total protein was used as a normalization method.

---

**Figure 1—Figure Supplement 1.**
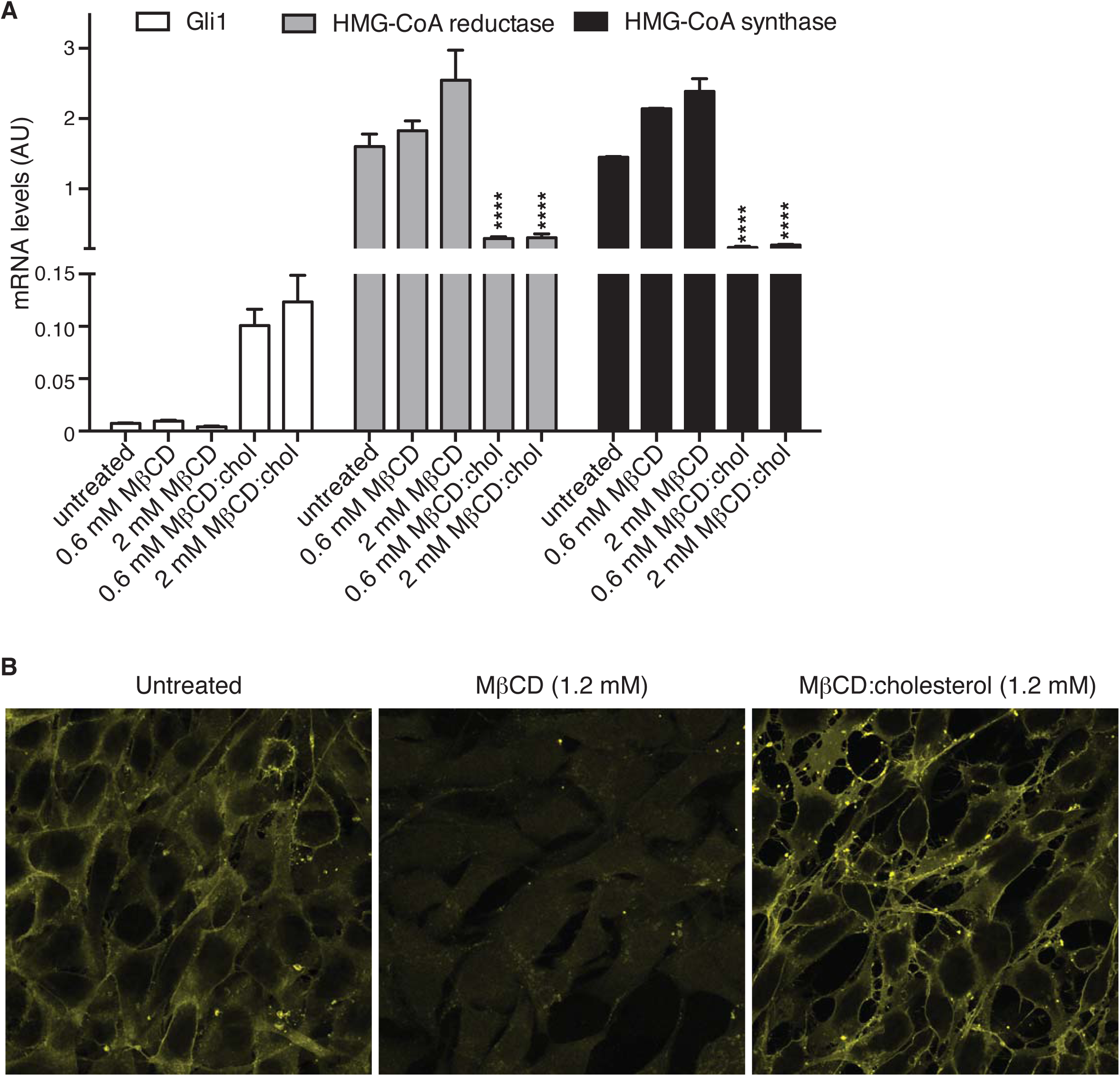
MβCD:cholesterol treatment increases the free cholesterol content of NIH/3T3 cells. (**A**) Mean (+/-SD, n=4) mRNA levels of *Gli1* or of two genes, encoding HMG-CoA reductase and synthase, in the cholesterol biosynthetic pathway that are negatively regulated by cellular cholesterol levels are shown after treatment with the indicated concentrations of MβCD or the MβCD:cholesterol complex. Asterisks denote statistical significance for difference from the untreated sample using two-way ANOVA with a Holm-Sidak post-test. (**B**) Levels of free cholesterol in the plasma membrane were assessed by staining with fluorescently labeled Perfringolysin O (PFO), a toxin that preferentially binds to the accessible (or chemically active) pool of cholesterol in membranes.

**Figure 3—Figure Supplement 1.**
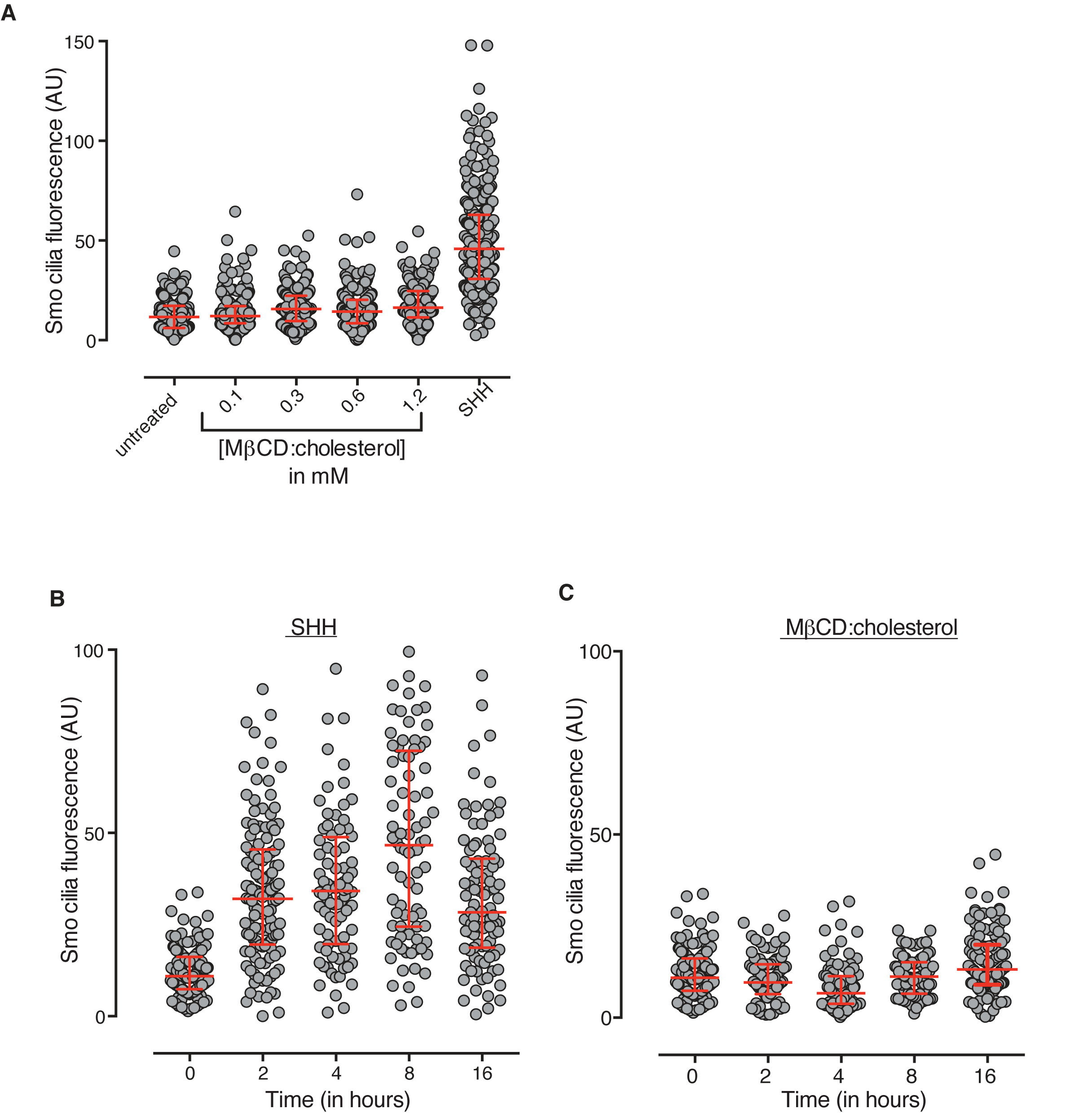
MβCD:cholesterol fails to drive SMO accumulation in the ciliary membrane. (**A**) SMO protein levels in primary cilia were determined by immunostaining NIH/3T3 cells after treatment (12 hours) with SHH (265 nM) or the indicated concentrations of MβCD:cholesterol. The kinetics of SMO accumulation in cilia were measured after treatment with SHH (**B**, 265 nM) or MβCD:cholesterol (**C**, 1.2 mM). Each point depicts SMO fluorescence at a single cilium and the red bars show the median and interquartile range of measurements from ∼250 cilia per condition for **A** and ∼100 cilia per condition for **B** and **C**.

**Figure 4—Figure Supplement 1.**
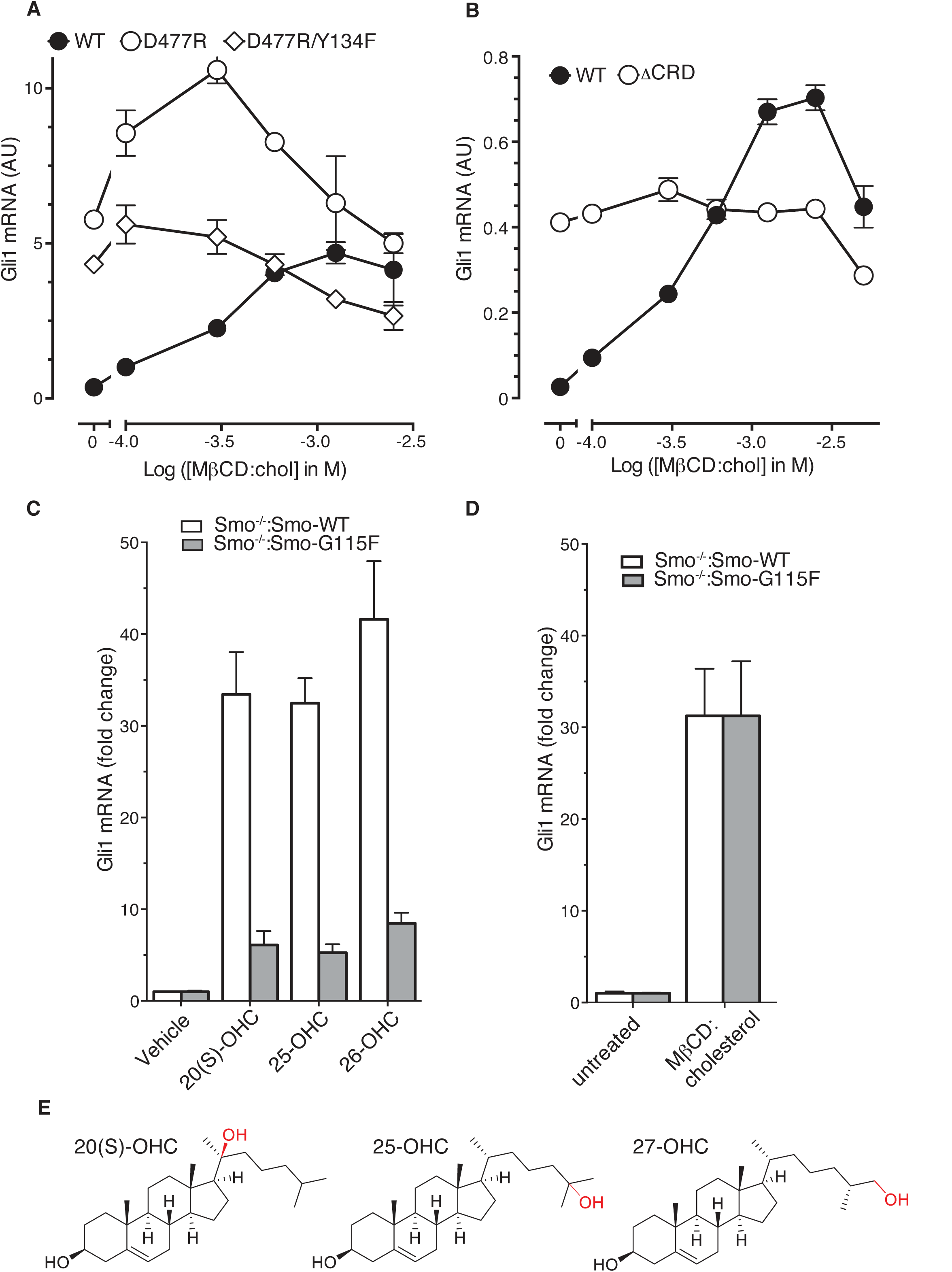
Role of the cysteine-rich domain of Smo in responses to cholesterol and side-chain oxysterols. (**A and B**) Dose-response curves for MβCD:cholesterol in Smo^−/−^ cells stably expressing WT SMO (solid black circles) or the indicated SMO mutants. D477R (**A**) is an activating mutation in the 7TMD, Y134F (**A**) is a mutation in the CRD (see Figure 5) that abrogates cholesterol and oxysterol responses, and ΔCRD (**B**) is an activating N-terminal truncation mutant that lacks the entire CRD. (**C**) *Gli1* induction in Smo^−/−^ cells expressing SMO-WT or SMO-G115F (see Figure 4 and associated discussion) treated with the indicated side-chain oxysterols, each applied at 5 μM as an inclusion complex with 44 μM MβCD. The activity of MβCD:cholesterol (1.2 mM) in both cell lines is shown in (**D**) for comparison. (**E**) Structures of the various side-chain oxysterols used in **C**, with differences from cholesterol highlighted in red.

## References

Bazan, J.F., and de Sauvage, F.J. (2009). Structural ties between cholesterol transport and morphogen signaling. Cell 138, 1055–1056 10.1016/j.cell.2009.09.006

Bidet, M., Joubert, O., Lacombe, B., Ciantar, M., Nehme, R., Mollat, P., Bretillon, L., Faure, H., Bittman, R., Ruat, M., et al. (2011). The hedgehog receptor patched is involved in cholesterol transport. PloS one 6, e23834 10.1371/journal.pone.0023834

Bishop, B., Aricescu, A.R., Harlos, K., O'Callaghan, C.A., Jones, E.Y., and Siebold, C. (2009). Structural insights into hedgehog ligand sequestration by the human hedgehog-interacting protein HHIP. Nat Struct Mol Biol 16, 698–703 10.1038/nsmb.1607

Blassberg, R., Macrae, J.I., Briscoe, J., and Jacob, J. (2016). Reduced cholesterol levels impair Smoothened activation in Smith-Lemli-Opitz syndrome. Human molecular genetics 25, 693–705 10.1093/hmg/ddv507

Briscoe, J., and Therond, P.P. (2013). The mechanisms of Hedgehog signalling and its roles in development and disease. Nat Rev Mol Cell Biol 14, 416–429 10.1038/nrm3598

Brown, A.J., Sun, L., Feramisco, J.D., Brown, M.S., and Goldstein, J.L. (2002). Cholesterol addition to ER membranes alters conformation of SCAP, the SREBP escort protein that regulates cholesterol metabolism. Mol Cell 10, 237–245

Brown, M.S., and Goldstein, J.L. (2009). Cholesterol feedback: from Schoenheimer's bottle to Scap's MELADL. J Lipid Res 50 Suppl, S15–27 10.1194/jlr.R800054-JLR200

Burger, K., Gimpl, G., and Fahrenholz, F. (2000). Regulation of receptor function by cholesterol. Cell Mol Life Sci 57, 1577–1592

Byrne, E.F., Sircar, R., Miller, P.S., Hedger, G., Luchetti, G., Nachtergaele, S., Tully, M.D., Mydock-McGrane, L., Covey, D.F., Rambo, R.P., et al. (2016). Structural basis of Smoothened regulation by its extracellular domains. Nature 535, 517–522 10.1038/nature18934

Carstea, E.D., Morris, J.A., Coleman, K.G., Loftus, S.K., Zhang, D., Cummings, C., Gu, J., Rosenfeld, M.A., Pavan, W.J., Krizman, D.B., et al. (1997). Niemann-Pick C1 disease gene: homology to mediators of cholesterol homeostasis. Science 277, 228–231

Chen, J.K., Taipale, J., Cooper, M.K., and Beachy, P.A. (2002a). Inhibition of Hedgehog signaling by direct binding of cyclopamine to Smoothened. Genes & development 16, 2743–2748

Chen, J.K., Taipale, J., Young, K.E., Maiti, T., and Beachy, P.A. (2002b). Small molecule modulation of Smoothened activity. Proc Natl Acad Sci U S A 99, 14071–14076

Cherezov, V., Rosenbaum, D.M., Hanson, M.A., Rasmussen, S.G., Thian, F.S., Kobilka, T.S., Choi, H.J., Kuhn, P., Weis, W.I., Kobilka, B.K., et al. (2007). High-resolution crystal structure of an engineered human beta2-adrenergic G protein-coupled receptor. Science 318, 1258–1265 10.1126/science.1150577

Christian, A.E., Haynes, M.P., Phillips, M.C., and Rothblat, G.H. (1997). Use of cyclodextrins for manipulating cellular cholesterol content. J Lipid Res 38, 2264–2272

Cooper, M.K., Porter, J.A., Young, K.E., and Beachy, P.A. (1998). Teratogen-mediated inhibition of target tissue response to Shh signaling. Science 280, 1603–1607

Cooper, M.K., Wassif, C.A., Krakowiak, P.A., Taipale, J., Gong, R., Kelley, R.I., Porter, F.D., and Beachy, P.A. (2003). A defective response to Hedgehog signaling in disorders of cholesterol biosynthesis. Nature genetics 33, 508–513

Corcoran, R.B., and Scott, M.P. (2006). Oxysterols stimulate Sonic hedgehog signal transduction and proliferation of medulloblastoma cells. Proc Natl Acad Sci U S A 103, 8408–8413

Covey, D.F. (2009). ent-Steroids: novel tools for studies of signaling pathways. Steroids 74, 577–585

Das, A., Goldstein, J.L., Anderson, D.D., Brown, M.S., and Radhakrishnan, A. (2013). Use of mutant 125I-perfringolysin O to probe transport and organization of cholesterol in membranes of animal cells. Proc Natl Acad Sci U S A 110, 10580–10585 10.1073/pnas.1309273110

Dessaud, E., McMahon, A.P., and Briscoe, J. (2008). Pattern formation in the vertebrate neural tube: a sonic hedgehog morphogen-regulated transcriptional network. Development 135, 2489–2503 10.1242/dev.009324

Dijkgraaf, G.J., Alicke, B., Weinmann, L., Januario, T., West, K., Modrusan, Z., Burdick, D., Goldsmith, R., Robarge, K., Sutherlin, D., et al. (2011). Small molecule inhibition of GDC-0449 refractory smoothened mutants and downstream mechanisms of drug resistance. Cancer research 71, 435–444 10.1158/0008-5472.CAN-10-2876

Dwyer, J.R., Sever, N., Carlson, M., Nelson, S.F., Beachy, P.A., and Parhami, F. (2007). Oxysterols are novel activators of the hedgehog signaling pathway in pluripotent mesenchymal cells. J Biol Chem 282, 8959–8968

Eaton, S. (2008). Multiple roles for lipids in the Hedgehog signalling pathway. Nat Rev Mol Cell Biol 9, 437–445 10.1038/nrm2414

Fahrenholz, F., Klein, U., and Gimpl, G. (1995). Conversion of the myometrial oxytocin receptor from low to high affinity state by cholesterol. Adv Exp Med Biol 395, 311–319

Frank-Kamenetsky, M., Zhang, X.M., Bottega, S., Guicherit, O., Wichterle, H., Dudek, H., Bumcrot, D., Wang, F.Y., Jones, S., Shulok, J., et al. (2002). Small-molecule modulators of Hedgehog signaling: identification and characterization of Smoothened agonists and antagonists. J Biol 1, 10

Gimpl, G., Burger, K., and Fahrenholz, F. (1997). Cholesterol as modulator of receptor function. Biochemistry 36, 10959–10974 10.1021/bi963138w

Gimpl, G., and Fahrenholz, F. (2002). Cholesterol as stabilizer of the oxytocin receptor. Biochim Biophys Acta 1564, 384–392

Gouti, M., Tsakiridis, A., Wymeersch, F.J., Huang, Y., Kleinjung, J., Wilson, V., and Briscoe, J. (2014). In vitro generation of neuromesodermal progenitors reveals distinct roles for wnt signalling in the specification of spinal cord and paraxial mesoderm identity. PLoS biology 12, e1001937 10.1371/journal.pbio.1001937

Haberland, M.E., and Reynolds, J.A. (1973). Self-association of cholesterol in aqueous solution. Proc Natl Acad Sci U S A 70, 2313–2316

Incardona, J.P., and Eaton, S. (2000). Cholesterol in signal transduction. Curr Opin Cell Biol 12, 193–203

Incardona, J.P., Gaffield, W., Kapur, R.P., and Roelink, H. (1998). The teratogenic Veratrum alkaloid cyclopamine inhibits sonic hedgehog signal transduction. Development 125, 3553–3562

Incardona, J.P., Gruenberg, J., and Roelink, H. (2002). Sonic hedgehog induces the segregation of patched and smoothened in endosomes. Current biology : CB 12, 983–995

Incardona, J.P., and Roelink, H. (2000). The role of cholesterol in Shh signaling and teratogen-induced holoprosencephaly. Cell Mol Life Sci 57, 1709–1719

Janda, C.Y., Waghray, D., Levin, A.M., Thomas, C., and Garcia, K.C. (2012). Structural basis of Wnt recognition by Frizzled. Science 337, 59–64 10.1126/science.1222879

Jessell, T.M. (2000). Neuronal specification in the spinal cord: inductive signals and transcriptional codes. Nature reviews Genetics 1, 20–29 10.1038/35049541

Jiang, X., and Covey, D.F. (2002). Total synthesis of ent-cholesterol via a steroid C,D-ring side-chain synthon. J Org Chem 67, 4893–4900

Khaliullina, H., Bilgin, M., Sampaio, J.L., Shevchenko, A., and Eaton, S. (2015). Endocannabinoids are conserved inhibitors of the Hedgehog pathway. Proc Natl Acad Sci U S A 112, 3415–3420 10.1073/pnas.1416463112

Khaliullina, H., Panakova, D., Eugster, C., Riedel, F., Carvalho, M., and Eaton, S. (2009). Patched regulates Smoothened trafficking using lipoprotein-derived lipids. Development 136, 4111–4121 10.1242/dev.041392

Klein, U., Gimpl, G., and Fahrenholz, F. (1995). Alteration of the myometrial plasma membrane cholesterol content with beta-cyclodextrin modulates the binding affinity of the oxytocin receptor. Biochemistry 34, 13784–13793

Kutejova, E., Sasai, N., Shah, A., Gouti, M., and Briscoe, J. (2016). Neural Progenitors Adopt Specific Identities by Directly Repressing All Alternative Progenitor Transcriptional Programs. Developmental cell 36, 639–653 10.1016/j.devcel.2016.02.013

Li, J., Deffieu, M.S., Lee, P.L., Saha, P., and Pfeffer, S.R. (2015). Glycosylation inhibition reduces cholesterol accumulation in NPC1 protein-deficient cells. Proc Natl Acad Sci U S A 112, 14876–14881 10.1073/pnas.1520490112

Lingwood, D., and Simons, K. (2010). Lipid rafts as a membrane-organizing principle. Science 327, 46–50 10.1126/science.1174621

Lopez, C.A., de Vries, A.H., and Marrink, S.J. (2011). Molecular mechanism of cyclodextrin mediated cholesterol extraction. PLoS Comput Biol 7, e1002020 10.1371/journal.pcbi.1002020

Myers, B.R., Sever, N., Chong, Y.C., Kim, J., Belani, J.D., Rychnovsky, S., Bazan, J.F., and Beachy, P.A. (2013). Hedgehog pathway modulation by multiple lipid binding sites on the smoothened effector of signal response. Developmental cell 26, 346–357 10.1016/j.devcel.2013.07.015

Nachtergaele, S., Mydock, L.K., Krishnan, K., Rammohan, J., Schlesinger, P.H., Covey, D.F., and Rohatgi, R. (2012). Oxysterols are allosteric activators of the oncoprotein Smoothened. Nature chemical biology 8, 211–220 10.1038/nchembio.765

Nachtergaele, S., Whalen, D.M., Mydock, L.K., Zhao, Z., Malinauskas, T., Krishnan, K., Ingham, P.W., Covey, D.F., Siebold, C., and Rohatgi, R. (2013). Structure and function of the Smoothened extracellular domain in vertebrate Hedgehog signaling. Elife 2, e01340 10.7554/eLife.01340

Nedelcu, D., Liu, J., Xu, Y., Jao, C., and Salic, A. (2013). Oxysterol binding to the extracellular domain of Smoothened in Hedgehog signaling. Nature chemical biology 9, 557–564 10.1038/nchembio.1290

Niewiadomski, P., Kong, J.H., Ahrends, R., Ma, Y., Humke, E.W., Khan, S., Teruel, M.N., Novitch, B.G., and Rohatgi, R. (2014). Gli protein activity is controlled by multisite phosphorylation in vertebrate hedgehog signaling. Cell Rep 6, 168–181 10.1016/j.celrep.2013.12.003

Novitch, B.G., Chen, A.I., and Jessell, T.M. (2001). Coordinate regulation of motor neuron subtype identity and pan-neuronal properties by the bHLH repressor Olig2. Neuron 31, 773–789

Pear, W.S., Nolan, G.P., Scott, M.L., and Baltimore, D. (1993). Production of high-titer helper-free retroviruses by transient transfection. Proc Natl Acad Sci U S A 90, 8392–8396

Porter, J.A., Young, K.E., and Beachy, P.A. (1996). Cholesterol modification of hedgehog signaling proteins in animal development. Science 274, 255–259

Prasanna, X., Chattopadhyay, A., and Sengupta, D. (2014). Cholesterol modulates the dimer interface of the beta(2)-adrenergic receptor via cholesterol occupancy sites. Biophys J 106, 1290–1300 10.1016/j.bpj.2014.02.002

Pucadyil, T.J., and Chattopadhyay, A. (2004). Cholesterol modulates ligand binding and G-protein coupling to serotonin(1A) receptors from bovine hippocampus. Biochim Biophys Acta 1663, 188–200 10.1016/j.bbamem.2004.03.010

Pucadyil, T.J., and Chattopadhyay, A. (2006). Role of cholesterol in the function and organization of G-protein coupled receptors. Prog Lipid Res 45, 295–333 10.1016/j.plipres.2006.02.002

Pusapati, G.V., Hughes, C.E., Dorn, K.V., Zhang, D., Sugianto, P., Aravind, L., and Rohatgi, R. (2014). EFCAB7 and IQCE regulate hedgehog signaling by tethering the EVC-EVC2 complex to the base of primary cilia. Developmental cell 28, 483–496 10.1016/j.devcel.2014.01.021

Radhakrishnan, A., and McConnell, H.M. (2000). Chemical activity of cholesterol in membranes. Biochemistry 39, 8119–8124

Rohatgi, R., Milenkovic, L., Corcoran, R.B., and Scott, M.P. (2009). Hedgehog signal transduction by Smoothened: pharmacologic evidence for a 2-step activation process. Proc Natl Acad Sci U S A 106, 3196–3201 10.1073/pnas.0813373106

Rohatgi, R., Milenkovic, L., and Scott, M.P. (2007). Patched1 regulates hedgehog signaling at the primary cilium. Science 317, 372–376 10.1126/science.1139740

Ruprecht, J.J., Mielke, T., Vogel, R., Villa, C., and Schertler, G.F. (2004). Electron crystallography reveals the structure of metarhodopsin I. The EMBO journal 23, 3609–3620 10.1038/sj.emboj.7600374

Sever, N., Mann, R.K., Xu, L., Snell, W.J., Hernandez-Lara, C.I., Porter, N.A., and Beachy, P.A. (2016). Endogenous B-ring oxysterols inhibit the Hedgehog component Smoothened in a manner distinct from cyclopamine or side-chain oxysterols. Proc Natl Acad Sci U S A 113, 5904–5909 10.1073/pnas.1604984113

Sharpe, H.J., Wang, W., Hannoush, R.N., and de Sauvage, F.J. (2015). Regulation of the oncoprotein Smoothened by small molecules. Nature chemical biology 11, 246–255 10.1038/nchembio.1776

Swamy, M., Beck-Garcia, K., Beck-Garcia, E., Hartl, F.A., Morath, A., Yousefi, O.S., Dopfer, E.P., Molnar, E., Schulze, A.K., Blanco, R., et al. (2016). A Cholesterol-Based Allostery Model of T Cell Receptor Phosphorylation. Immunity 44, 1091–1101 10.1016/j.immuni.2016.04.011

Taipale, J., Cooper, M.K., Maiti, T., and Beachy, P.A. (2002). Patched acts catalytically to suppress the activity of Smoothened. Nature 418, 892–897

Theunissen, J.J., Jackson, R.L., Kempen, H.J., and Demel, R.A. (1986). Membrane properties of oxysterols. Interfacial orientation, influence on membrane permeability and redistribution between membranes. Biochim Biophys Acta 860, 66–74

Varjosalo, M., Li, S.P., and Taipale, J. (2006). Divergence of hedgehog signal transduction mechanism between Drosophila and mammals. Developmental cell 10, 177–186

Wang, C., Wu, H., Evron, T., Vardy, E., Han, G.W., Huang, X.P., Hufeisen, S.J., Mangano, T.J., Urban, D.J., Katritch, V., et al. (2014). Structural basis for Smoothened receptor modulation and chemoresistance to anticancer drugs. Nat Commun 5, 4355 10.1038/ncomms5355

Wichterle, H., Lieberam, I., Porter, J.A., and Jessell, T.M. (2002). Directed differentiation of embryonic stem cells into motor neurons. Cell 110, 385–397

Wu, H., Wang, C., Gregory, K.J., Han, G.W., Cho, H.P., Xia, Y., Niswender, C.M., Katritch, V., Meiler, J., Cherezov, V., et al. (2014). Structure of a class C GPCR metabotropic glutamate receptor 1 bound to an allosteric modulator. Science 344, 58–64 10.1126/science.1249489

Yancey, P.G., Rodrigueza, W.V., Kilsdonk, E.P., Stoudt, G.W., Johnson, W.J., Phillips, M.C., and Rothblat, G.H. (1996). Cellular cholesterol efflux mediated by cyclodextrins. Demonstration Of kinetic pools and mechanism of efflux. J Biol Chem 271, 16026–16034

Yauch, R.L., Dijkgraaf, G.J., Alicke, B., Januario, T., Ahn, C.P., Holcomb, T., Pujara, K., Stinson, J., Callahan, C.A., Tang, T., et al. (2009). Smoothened mutation confers resistance to a Hedgehog pathway inhibitor in medulloblastoma. Science 326, 572–574

Ying, Q.L., Stavridis, M., Griffiths, D., Li, M., and Smith, A. (2003). Conversion of embryonic stem cells into neuroectodermal precursors in adherent monoculture. Nat Biotechnol 21, 183–186 10.1038/nbt780

Zidovetzki, R., and Levitan, I. (2007). Use of cyclodextrins to manipulate plasma membrane cholesterol content: evidence, misconceptions and control strategies. Biochim Biophys Acta 1768, 1311–1324 10.1016/j.bbamem.2007.03.026

